# Learning sequence, structure, and function representations of proteins with language models

**DOI:** 10.1101/2023.11.26.568742

**Authors:** Tymor Hamamsy, Meet Barot, James T. Morton, Martin Steinegger, Richard Bonneau, Kyunghyun Cho

**Affiliations:** Center for Data Science, New York University, New York, NY, USA; Mythos Scientific, NJ, USA; Biostatistics & Bioinformatics Branch, Eunice Kennedy Shriver National Institute of Child Health and Human Development, National Institutes of Health, Bethesda, MD, USA; School of Biological Sciences, Seoul National University, Seoul, South Korea; Artificial Intelligence Institute, Seoul National University, Seoul, South Korea; Institute of Molecular Biology and Genetics, Seoul National University, Seoul, South Korea; Prescient Design, gRED Computational Sciences, Genentech, New York, NY, USA; CIFAR Fellow; Department of Computer Science, Courant Institute of Mathematical Sciences, New York University, New York, NY, USA

## Abstract

The sequence-structure-function relationships that ultimately generate the diversity of extant observed proteins is complex, as proteins bridge the gap between multiple informational and physical scales involved in nearly all cellular processes. One limitation of existing protein annotation databases such as UniProt is that less than 1% of proteins have experimentally verified functions, and computational methods are needed to fill in the missing information. Here, we demonstrate that a multi-aspect framework based on protein language models can learn sequence-structure-function representations of amino acid sequences, and can provide the foundation for sensitive sequence-structure-function aware protein sequence search and annotation. Based on this model, we introduce a multi-aspect information retrieval system for proteins, Protein-Vec, covering sequence, structure, and function aspects, that enables computational protein annotation and function prediction at tree-of-life scales.

## Introduction

The sequence-structure-function relationship is central to biology; it states that a protein’s amino-acid sequence determines its 3D structure (or ensemble of structures), which in turn determines its biological functions [1, 2, 3, 4]. The relationships between sequence, structure and function can be extraordinarily complex, and must incorporate and account for evolution, structural dynamics and neo-functionalization of genes in different organismal and cellular contexts (each example complexity here warrents the founding of a proper biological sub-field). In spite of this daunting complexity, disentangling this relationship is a worthwhile goal, as understanding and explaining relationships across diverse sequences can uncover novel biological insights. Annotating and navigating vast sequence databases is also key to understanding biodiversity (where large sequence databases are a main way we record biodiversity, currently our most valuable planetary export). In this work, we introduce a new multi-aspect information retrieval system for proteins, covering sequence, structure, and function aspects, that enables multi-aspect retrieval at tree-of-life scales.

At the time of this writing, there are over 250 million proteins in UniProt [5], there are only around 570K proteins that have been reviewed by humans in Swiss-Prot, and an even fewer number of these proteins have been experimentally validated [5]. As over one third of proteins cannot be functionally annotated by sequence alignment methods [6, 7], computational protein annotation and function prediction have become essential to protein annotation. Large numbers of methods for gene and protein-gene annotation rely on mapping annotation (or subsets of annotations) via inference/detection of homology, and methods for remote homology detection remain the most used tools in bioinformatics to this date. Several lines of thinking can be used to estimate the fraction of unannotated genes that are unique (on origin or function) vs. the number of proteins that seem unique only due to a lack of homology-detection sensitivity. Studies of neutral rates of evolution as well as studies of protein-protein structure similarity (and structure similarity correspondence with function concordance), suggest that a large fraction of novel or unannotated proteins are cases of failed homology detection [8, 9, 10]. Protein multi-aspect function prediction and associated homology detection problems remain a top priority in computational biology, for example several methods exist to perform protein function prediction and annotation that also include a wide range of data beyond sequence [11, 12, 13, 14, 15, 16, 17, 18, 19, 20, 21, 22, 23, 24, 25], including methods that use protein sequences [15, 18, 19, 21], protein-protein interaction networks [15, 26, 27], protein structures [17], and metabolomics [28, 29, 30]

Here, we approach computational protein annotation and function prediction by modeling different protein aspects and in the process, building a multi-aspect information retrieval system. The different aspects that we model include protein function (encapsulated in Enzyme Commission numbers (EC) and Gene Ontology (GO) annotations [31]), protein domain sequence families (Pfam) [12], gene domains (Gene3D) [32], and protein structure information (via TM-Scores) [33]. Each aspect has its own unique computational and completeness challenges. For EC classification, while there have been many computational tools developed to characterize enzyme function [5, 13, 34, 35, 36, 37], a recent study found that approximately 40% of computationally annotated enzymes are incorrectly annotated [19]. While Pfam consists of protein domain sets that were manually curated, many Pfam families consist of small sets of sequences [38], complicating or preventing training for many machine learning approaches. Protein function prediction (GO) has historically been challenged by incorrect GO annotations [39], sparse experimental GO annotations [40], and issues relating to it being an extreme multi-label multi-class classification problem. Protein function prediction from sequences alone is perhaps the most common modeling approach as sequences are the most widely accessible modality, and double-blind tests of methods, such as CAFA, have emerged to benchmark and compare different protein function methods that use sequences alone [21].

In this work, we introduce Protein-Vec, a deep learning model for multi-aspect information retrieval (IR). Vector-based information retrieval systems for proteins have recently achieved excellent results for structure-aware search [41]. As proteins have many important aspects across sequence, structure, and function, it is valuable to design an IR system for proteins that can generate meaningful vector representations of proteins for different aspects and combinations of aspects. Altogether, we propose a full information retrieval pipeline that can compete with state-of-the-art sequence and structure search + alignment approaches as well as function prediction methods. The aspects that we chose to study are central to the sequence-structure-function of proteins, but the approach we have introduced is flexible and in future work we plan to train a model on even more relevant protein aspects. Our approach integrates these aspects into informative protein features that allow for state-of-the-art performances across all individual aspects’ evaluation tasks.

## Results

Our contribution is to introduce an approach to perform scalable sequence-structure-function aware protein search, Protein-Vec (Figures 1, 2, 3), and in the process, we introduce a framework to perform multi-aspect information retrieval. Our overall approach involves contrastive learning [42], the conditional generation of protein vectors based on the aspects being queried, and deep neural networks (Figures 1, 3). Once a Protein-Vec model is trained, it can produce multi-aspect-aware vectors for proteins that can be indexed and queried (Figure 2).

**Figure 1:**
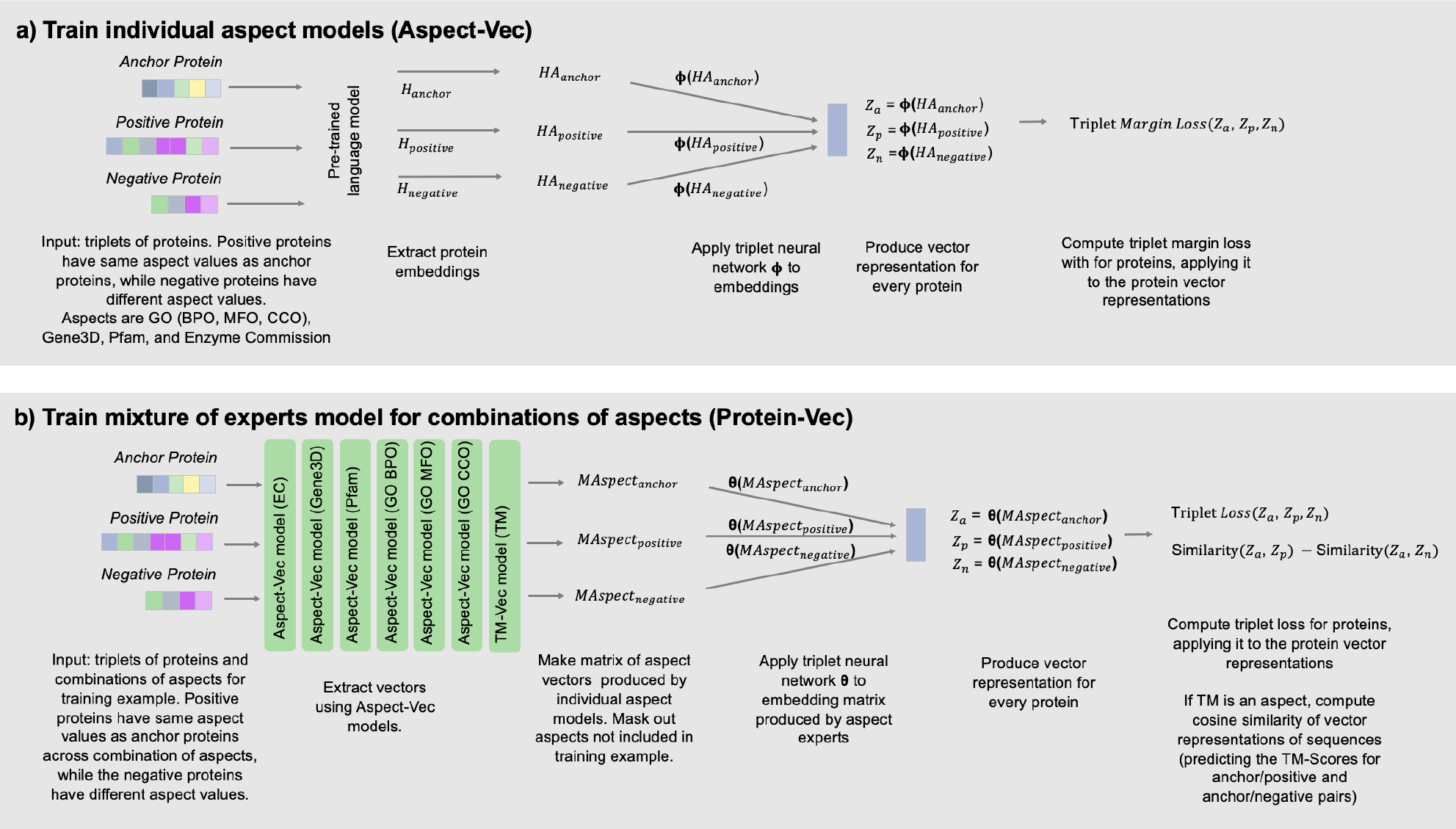
Overview of Protein-Vec training pipeline. Overview of Protein-Vec training pipeline. Training a Protein-Vec model consists of two stages. First, we trained individual Aspect-Vec models for the different protein aspects we consider, including TM-Scores, GO-MFO, GO-BPO, GO-CCO, EC, Gene3D, and Pfam. Then, we train a mixture of experts model, that is built on top of all of the Aspect-Vec models (experts in this case). a) Each Aspect-Vec model (besides the TM-Score Aspect-Vec model, which is simply a fully trained TM-Vec model) is trained on triplets of amino acid sequences. Each triplet, consists of an anchor protein, a positive example (a protein that shares a label with the anchor protein), and a negative example (a protein that has a different label than the anchor protein). When we input a triplet of sequences (i.e. domains, chains, proteins), we first apply a pretrained deep protein language model to extract embeddings for every residue of each sequence. Then, we apply a triplet neural network, called *ϕ*, to the embeddings of each sequence, and produce a vector representation, *z*, for each sequence. The *ϕ* network is trained on millions of triplets of sequences for every aspect that we build an Aspect-Vec model for. Finally, we compute the triplet margin loss of the vector representations. b) At the second stage of training, we build a mixture of experts model, called Protein-Vec. In every training example, we input triplets of proteins, as well a combinations of aspects for the training example. In every triplet, positive examples share the same aspect values as the anchor protein across all of the aspects considered in the training example, and negative examples have different aspect values from the anchor protein across the considered aspects. Next, we apply the 7 Aspect-Vec models (EC, Gene3D, Pfam, GO-BPO, GO-MFO, GO-CCO, and TM) to each sequence in the triplet. We are now left with a matrix of aspect vectors for each triplet, of size 7 (number of aspects) by 512 (vector dimension for each aspect). For a given training example, we mask out the aspects which are not specified in the training example, and we then apply a new triplet neural network to them, *θ*, which is our mixture of experts model. Finally, we compute a custom loss function, which has two components, a triplet margin loss component and a loss component for TM-Score predictions (if TM-Score is one of the training example’s aspects).

**Figure 2:**
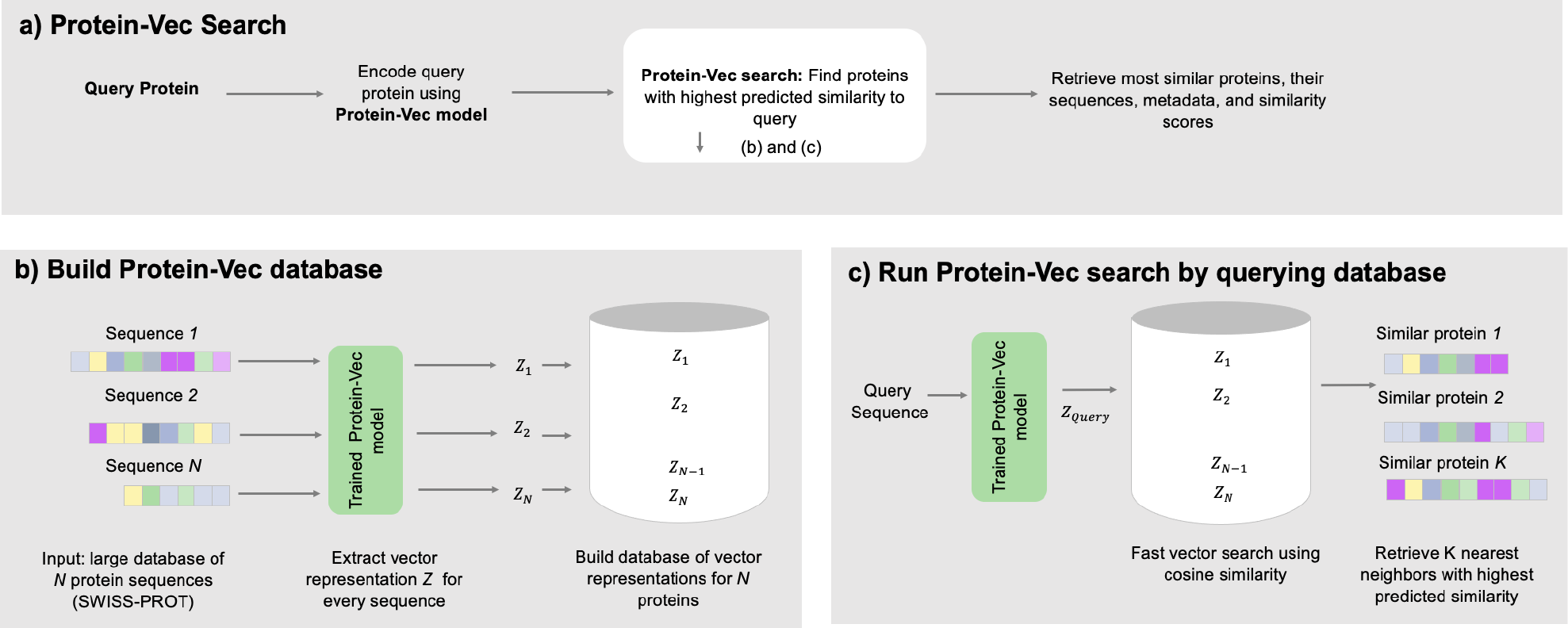
Overview of Protein-Vec search pipeline. (a) A Protein-Vec search pipeline consists of two stages: embedding and search. First, a query protein sequence is embedded using a Protein-Vec model. Then, Protein-Vec search is applied and nearest neighbor proteins are retrieved. These nearest neighbor proteins have the most similar Protein-Vec embeddings to the query protein’s Protein-Vec embeddings (reflecting sequence-structure-function aspects of proteins). (b) We build a Protein-Vec database by encoding large databases of protein sequences using a trained Protein-Vec model. For example, we inputted the protein sequences included in Swiss-Prot, extracted vector representations for every sequence, and built an indexed database of Protein-Vec’s sequence-structure-function aware vector representations of proteins. (c) Demonstration of Protein-Vec search using the Protein-Vec pipeline. After an indexed database of vector representations has been built, protein search consists of encoding the query sequence using a trained Protein-Vec model, and then performing fast vector search using cosine similarity as the search metric. Finally, we retrieve the K nearest neighbors with the highest predicted embedding similarity to the query proteins, as well as their annotations for different protein aspects. The predicted annotations for a given query protein come from the top-1 nearest neighbor protein’s annotations.

**Figure 3:**
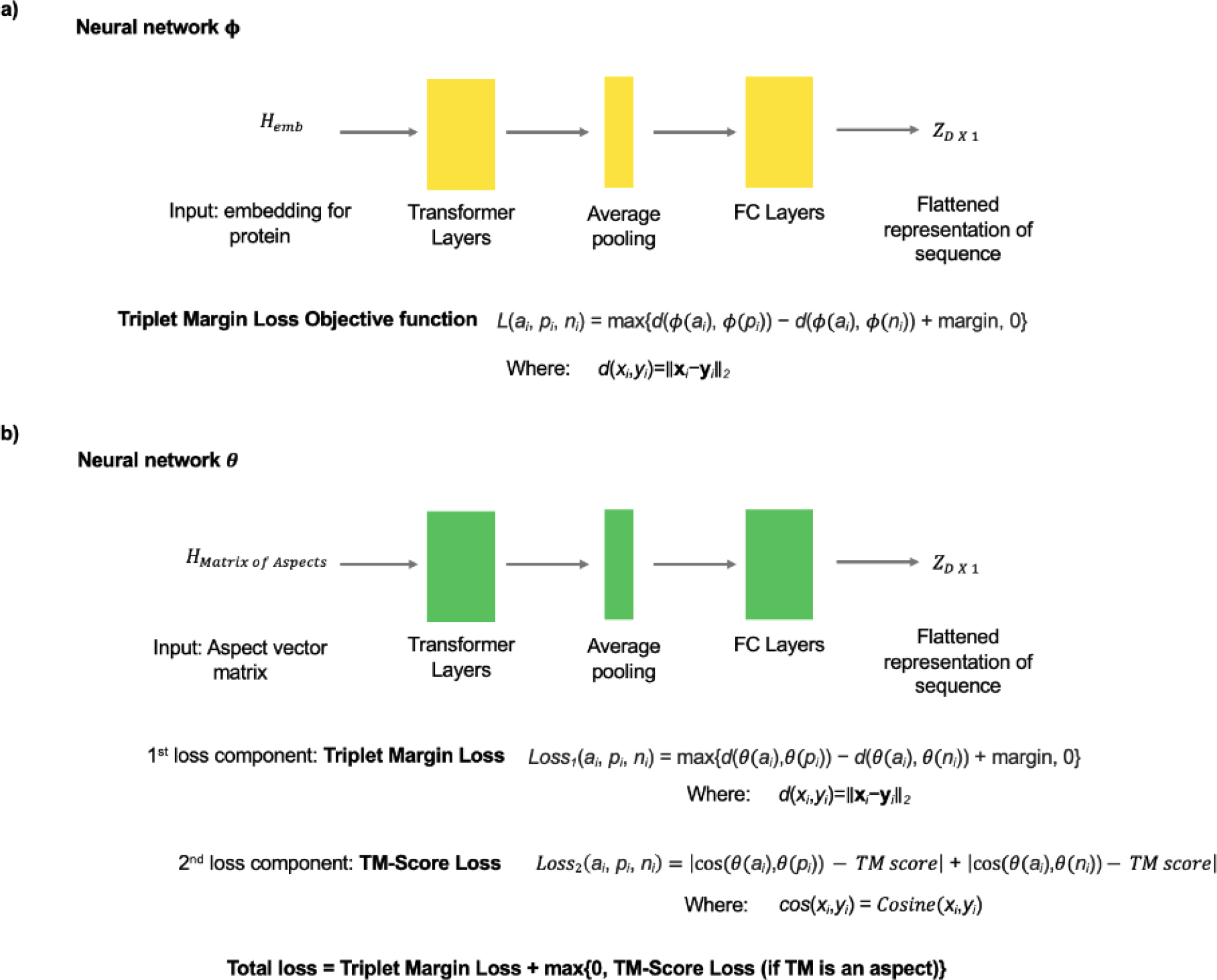
a) Overview of Aspect-Vec neural network architecture. The function *ϕ* takes in residue embeddings for each protein in the triplet and produces a flattened vector representation for each protein. *ϕ* is composed of several transformer encoder layers, followed by average pooling, dropout, and fully connected layers. At the final step, we calculate the triplet margin loss using the vector representations of each protein in the triplet as well as the margin, and our training objective is to minimize the triplet margin loss. b) Overview of Protein-Vec neural network architecture. The function *θ* takes in an aspect matrix for each protein in the triplet and produces a flattened vector representation for each protein. The loss function for Protein-Vec consists of two parts, a triplet margin loss, and a TM-Score loss component. The TM-Score loss component is applied when TM-Scores are included as an aspect in the training example.

We encode sequence, structure, and function information into these protein vectors, by training a mixture of experts based neural network [43] using contrastive learning [42] with a modified loss function (Figures 1, 3). Before training a Protein-Vec model, we must first train single aspect models, which we call Aspect-Vec models or experts. We train these models for individual protein aspects, including Enzyme Commission number (EC), Gene Ontology (GO) [31], Protein Family (Pfam) [12], TM-Scores [33], and Gene3D domain annotations [32]. In every training sample for both the Aspect-Vec and Protein-Vec models, there is an anchor protein, a positive protein, and a negative protein. One of the training objectives is then to identify the positive protein (shares a label with the anchor protein) and differentiate it from the negative protein (has a different label than the anchor protein).

Once the Aspect-Vec models are trained, they become “experts” that can be used in our multi-aspect model, Protein-Vec. As we describe in Figure 1, we leverage these “experts” by creating a matrix consisting of Aspect-Vec vectors for every training example (masking out aspects not specified) and applying a mixture of experts neural network to the resulting matrix. Therefore, for every aspect in a given training sample (i.e. some combination of protein aspects), we first encode the anchor, positive, and negative protein sequences using the relevant Aspect-Vec or expert models, and then we train a mixture of experts, Protein-Vec model (Figure 1). Protein-Vec is similarly trained to discern positive and negative samples across combinations of multiple protein aspects, including EC number, GO, Pfam, TM-Scores, and Gene3D domain annotations.

Once a Protein-Vec model is trained and a lookup database is encoded, it is possible to perform fast sequence-structure-function aware protein searches by identifying nearest neighbors within the embedding space of the resulting Protein-Vec vector embedding database (Figure 2). Therefore, for any protein sequence, a single forward pass of our model can predict a multi-aspect-aware vector of the protein sequence. We showcase the ability of Protein-Vec to encode sequence, structural, and functional information from protein sequences using different benchmarks across the protein aspects we investigated.

### Aspect-Vec: Single-aspect expert models

We include 7 different single-aspect experts, referred to as Aspect-Vec models, that are part of the overarching Protein-Vec model. These aspects include EC number, Gene Ontology molecular function (GO MFO), Gene Ontology biological process (GO BPO), Gene Ontology cellular component (GO CCO), Pfam, TM-Scores [41], and Gene3D domain annotations [12, 32, 31, 33]. Before detailing Protein-Vec’s results, we discuss the results of the individual Aspect-Vec models.

### Enzyme Commission number prediction for novel proteins

We benchmark Aspect-Vec against a recent enzyme commission number prediction method: CLEAN [34]. Our evaluation is a time-based evaluation, we predict enzyme commission numbers for held out proteins from UniProt that were introduced to the database after May 25th, 2022 (we only train this Aspect-Vec model on proteins deposited before April 2022). On this benchmark, an Aspect-Vec model trained on only EC triplets significantly outperforms CLEAN (Table 6). The Aspect-Vec model achieves an exact match accuracy of 54% while CLEAN [34] achieves an exact match accuracy 45%.

### Pfam prediction for remote homologs

Many proteins cannot be functionally annotated by sequence alignment approaches, and this annotation gap is especially large for remote homologs. Proteins are also composed of distinct folded and/or disordered domains, and these domains are observed in shuffled and rearranged to compose an even greater diversity of overarching organismal functions. Thus, detection of very remote homology, and organization of protein segments into families is often centered on domains [44, 45]. In order to evaluate our ability to perform Pfam-centered remote homology detection using an Aspect-Vec model, we use a benchmark dataset constructed by clustering Pfam-seed v32 based on sequence identity where there is single-linkage clustering at 25% sequence identity within each Pfam family [46]. It is worth noting that this benchmark dataset is made up of protein domains whilst the queries/searches are whole proteins typically comprised of multiple domains. The test benchmark dataset (described in the Online Methods) consists of 21,293 distant held-out Pfam sequences. For assigning Pfam annotations to query (test) proteins, we assign the Pfam annotations of the nearest neighbor protein in the lookup (training) dataset to the query (test) proteins.

Table 1 shows that the Aspect-Vec model outperforms Top-Pick HMM [47], phmmer [47], BLASTp [48], ProtCNN [46], and ProtENN (which is an ensemble of 19 ProtCNN models) by wide margins [46]. Besides Aspect-Vec (Pfam), which has an error rate of 5.20%, the next best performing method is ProtENN at 12.20% on this benchmark.

**Table 1:** Comparing the performance of Aspect-Vec (Pfam) on a Pfam prediction benchmark. An Aspect-Vec model trained to predict Pfam annotations is compared against several different methods. The training dataset (which we use as our lookup dataset) and test datasets are comprised of protein domains, and were acquired from the ProtENN paper [46]. Every protein domain in the test dataset is at least 75% different in sequence similarity from every training protein domain. The Aspect-Vec (Pfam) Top 1 error rate represents the percentage of nearest neighbor protein domain’s Pfam annotations (our predictions) that are not identical to the test domain’s Pfam annotation.

| **Model** | **Error Rate (%)** |
| --- | --- |
| phmmer | 32.60% |
| BLASTp | 35.90% |
| ProtCNN | 27.60% |
| ProtENN | 12.20% |
| Aspect-Vec (Pfam) Top 1 | 5.20% |

### Predicting Gene-Ontology annotations for novel proteins

The Gene-Ontology is a widely used ontology to describe the functional properties of proteins, and predicting GO terms is a central bioinformatics task. Many protein function prediction competitions, such as CAFA, use time-based benchmarks to evaluate GO prediction methods. That is, a hidden, held-out dataset of newly experimentally annotated proteins is used to evaluate methods [21]. We use a time-based dataset from Swiss-Prot to compare Aspect-Vec against different function prediction methods, shown in Table 2 (see Online Methods). As most proteins have multiple GO annotations (often 10+) across the 3 GO categories (cellular component, biological process, and molecular function), our annotation procedure of assigning the GO annotations of the nearest neighbor proteins to query (test) proteins is a proof of concept that demonstrates the functional awareness of Aspect-Vec models.

**Table 2:** Comparing the performance of Aspect-Vec (GO) on a 2016 time-based split. The training and test datasets used for this benchmark come from the DeepGoPlus paper [16]. The F1 Max Scores of different GO prediction methods are compared on a held out dataset (2016 time-split).

| **Model** | **MFO** | **BPO** | **CCO** |
| --- | --- | --- | --- |
| Naïve | 31% | 32% | 61% |
| DiamondBLAST | 53% | 44% | 59% |
| DeepGO | 45% | 40% | 67% |
| DeepGOPlus | 59% | 47% | 70% |
| Aspect-Vec (GO) Top 1 | 54% | 41% | 64% |

On this time-based benchmark, we compare the F1 max scores of different Aspect-Vec models (GO-MFO, GO-BPO, and GO-CCO) against a naive method introduced in CAFA [21], DiamondBlast [49], DiamondScore [49], DeepGO [15], and DeepGOPlus [16]. Here we test a basic annotation pipeline of assigning all of the GO annotations of the nearest neighbor protein to query (test) proteins, and use a small lookup dataset (the training datasets in this benchmark are small for capturing the full advantages of nearest-neighbor lookup), our F1 max scores for the MFO and BPO aspects are higher than DeepGO’s (Table 2). The Aspect-Vec models achieve F1 max scores of 54%, 41%, and 64% on the GO-MFO, GO-BPO, and GO-CCO aspects, respectively. They are lower than but comparable to those by even the best performing method, DeepGOPlus [16], which achieves F1 max scores of 59%, 47%, and 70%.

### Predicting Gene3D domains: Diamond benchmark

Next, we build an Aspect-Vec model for Gene3D domains. Gene3D provides CATH annotations for the predicted domains in protein sequences [50]. Here, we evaluate the Aspect-Vec model’s performance on the DIAMOND benchmark [51] (see Online methods), which has UniRef50 [52] as a lookup database, and is comprised of both single and multiple-domain proteins. DIAMOND has a sensitivity of 99% for the top-1 proteins on this benchmark. After embedding proteins in the UniRef50 lookup database using an Aspect-Vec model trained on just Gene3D aspect annotations, we compare our performance on all query proteins up to 1,000 residues long. The nearest neighbor shares the same SCOP [53] family annotations 83% of the time (significantly lower than DIAMOND’s sensitivity of 99%). In comparison to the Gene3D Aspect-Vec model, TM-Vec [41] (which is instead trained to predict TM-Scores), achieves a much higher sensitivity on this benchmark of 92%.

### Protein-Vec: Combining sequence-structure-function aspects into one model

After evaluating the individual Aspect-Vec models, we study the performance of the proposed mixture-of-experts model, Protein-Vec.

In order to perform this analysis, we encode a Protein-Vec lookup database using proteins deposited to Swiss-Prot up to May 25, 2022. We then benchmark our Protein-Vec model on 3 different test datasets: a time-split test dataset of new proteins that were deposited to Swiss-Prot after May 25, 2022; a held-out test dataset where no proteins had more than 50% sequence similarity to the training proteins; and a held-out test dataset where no proteins had more than 20% sequence similarity to the training proteins. For every query protein, our predictions for the protein are the annotations of the nearest neighbor protein in our lookup dataset.

As the first step, we visualize Protein-Vec embeddings for held out proteins, coloring proteins by the different aspects (Figures 4, 5). Figure 4A shows a t-SNE plot of Protein-Vec embeddings for the held-out-20 and held-out-50 datasets, colored by their enzyme commission numbers. In both panels, there is a clear clustering by enzyme commission number, and a separation between different categories. As we use hard negative mining to find negative examples for triplets (see Online methods), we optimize for a distinction between EC classes at the 4-th and final level of the hierarchy (most granular). Figure 4B shows a t-SNE plot of Protein-Vec embeddings for the held-out-20 and held-out-50 datasets, colored by their Pfam annotations. In contrast to other Pfam annotation models that operate at the individual domain level (such as ProtENN), we can see in Figure 4B, that Protein-Vec can distinguish clusters for proteins with many Pfam annotations (i.e. clusters shown for PF10672; PF02926; PF01170 and PF02938; PF00152; PF01336). Figure 4C shows a t-SNE plot of Protein-Vec embeddings for the held-out-20 and held-out-50 datasets, colored by their Gene3D annotations. In this plot, we also notice protein clustering, even when proteins have many domains/Gene3D annotations. Across all of the plots in Figure 4, it is noteworthy that the clusters are tighter in the held-out-50 panels than in the held-out-20 panels; this is due to there being higher sequence similarity between the Protein-Vec training dataset and the held-out 50 dataset than between the training dataset and the held-out 20 dataset. Figure 5 shows a t-SNE plot for each of the GO components: MFO, BPO, and CCO. Across Figure 5A, Figure 5B, and Figure 5C, we see that Protein-Vec can successfully cluster proteins with multiple GO annotations. We again observe an improvement in the clustering in the held-out 50 panels compared to the held-out 20 panels.

**Figure 4:**
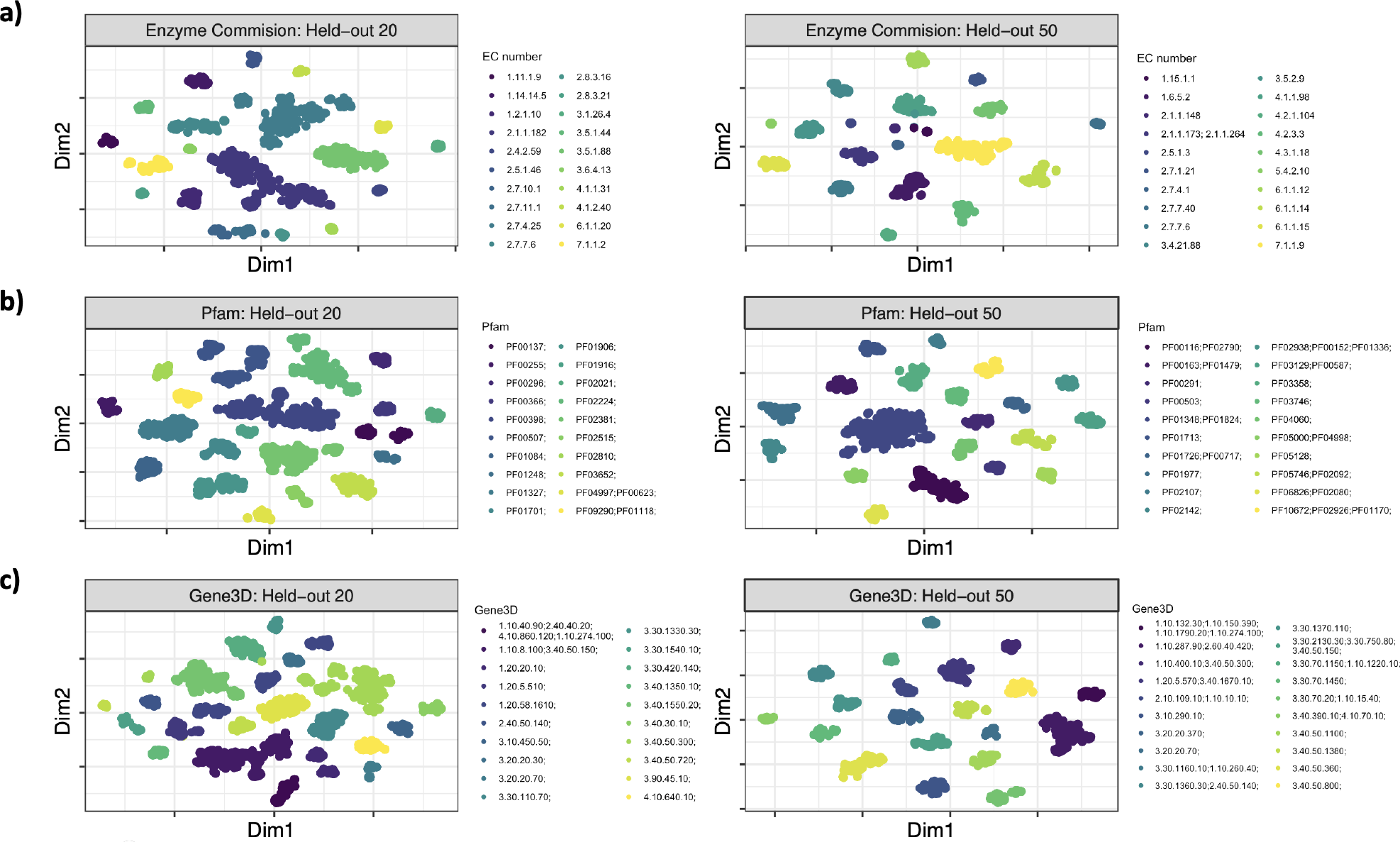
t-SNE plots of Protein-Vec embeddings of held out proteins at different sequence similarity thresholds. Here, we made t-SNE visualizations of the Protein-Vec embeddings for held out proteins; each plot shows a different aspect, which is colored by annotation values for that aspect. We only plotted the top 20 categories for each aspect. a) t-SNE plot of Protein-Vec embeddings, colored by Enzyme commission numbers. There is a clear separation of classes, and this separation improves from the held-out 20 dataset to the held-out 50 dataset. b) t-SNE plot of Protein-Vec embeddings, colored by Pfam annotations. c) t-SNE plot of Protein-Vec embeddings, colored by Gene3D annotations.

**Figure 5:**
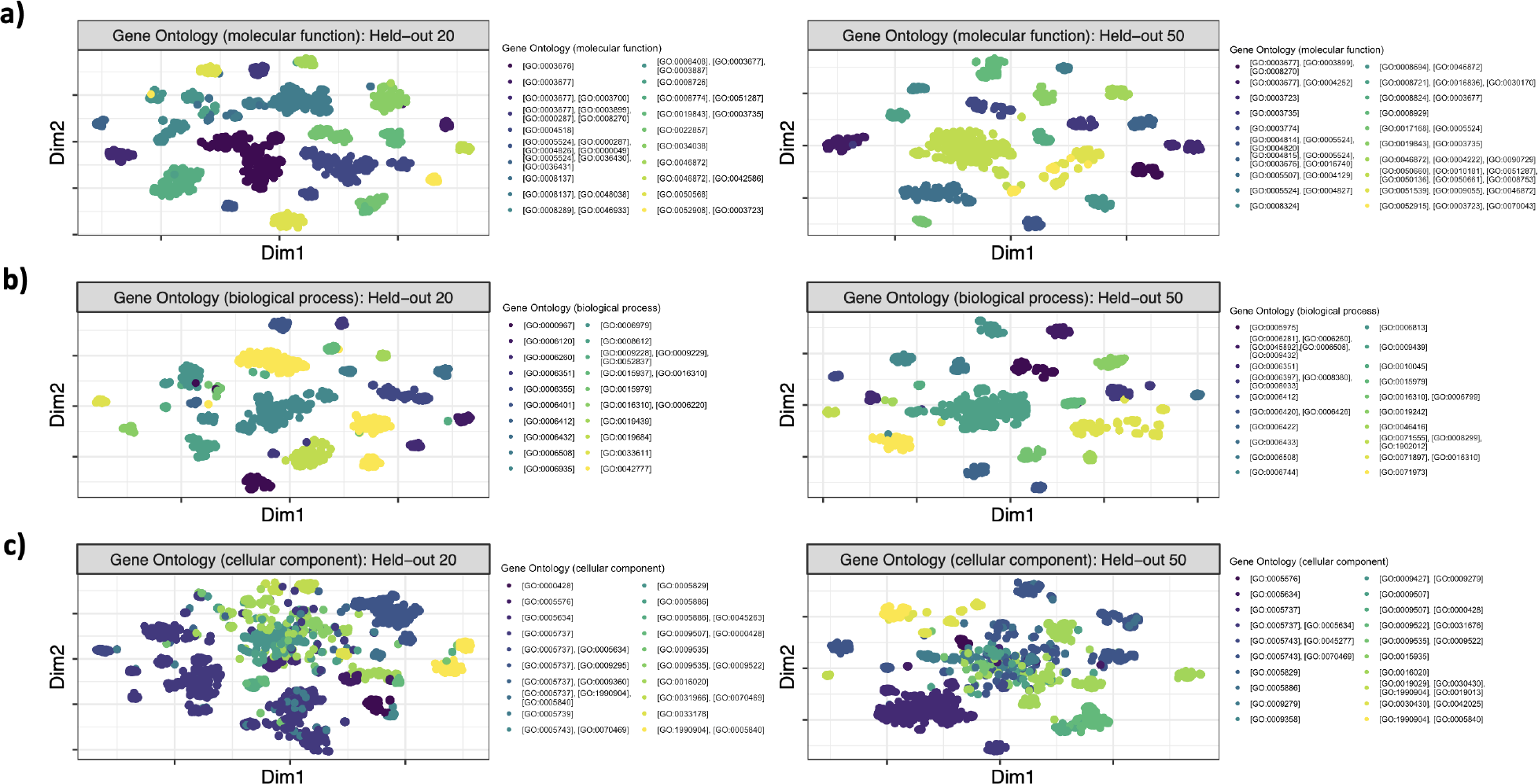
t-SNE plots of Protein-Vec embeddings of held out proteins at different sequence similarity thresholds, colored by their GO annotations. Here, we made t-SNE visualizations of the Protein-Vec embeddings for held out proteins; each plot shows a different GO aspect, which is colored by annotation values for that GO aspect. We only plotted the top 20 categories for each GO aspect. a) t-SNE plot of Protein-Vec embeddings, colored by GO-MFO annotations. b) t-SNE plot of Protein-Vec embeddings, colored by GO-BPO annotations. c) t-SNE plot of Protein-Vec embeddings, colored by GO-CCO annotations. Among the 3 GO aspects, GO-CCO has the least separation among its top categories, as can be seen in the held-out 50 and held-out 20 plots.

After visualizing the Protein-Vec embeddings of the held-out 50 and held-out 20 proteins, we next compare our multi-label, multi-classification performance on these test datasets (Table 3, Table 4). Table 3 shows the recall of Protein-Vec applied to the held-out 50% and held out 20% sequence similarity datasets. In Table 3, we define recall as the percentage of predicted annotations for an aspect that are among the ground truth annotations for the aspect. Across EC, Pfam, Gene3D, GO-MFO, GO-CCO, and GO-BPO, we note that the recall on the held-out 50% is higher than the recall on the held out 20%. This difference is largest for EC (where performance declines from 87% to 61%) and GO-CCO (where performance declines from 72% to 50%). As Table 3 demonstrates, our recall is extremely high for the held out 50% proteins, at 87%, 93%, 95% for EC, Pfam, and Gene3D respectively, and at 80%, 72%, and 75% for GO-MFO, GO-CCO, and GO-BPO respectively. We next calculate and report the exact match accuracy of Protein-Vec for the held out 50% and held out 20% datasets (Table 4). Here, an exact match implies that all of the predicted annotations for the aspect are identical to the ground truth annotations for the aspect. The exact match accuracy numbers for EC are nearly identical to the accuracy reported in Table 3. Pfam, and Gene3D performance both dropped in accuracy for the exact matches on the held-out 50% dataset from 93% to 88% (Pfam) and from 95% to 91% (Gene3D), accounting for the fact that many of the proteins in our held out datasets have multiple domains (and consequently, multiple Pfam and Gene3D annotations). Our performance predicting the exact GO annotations for held-out protein is notable; on the held-out 50% dataset, we correctly predict all of the GO annotations for proteins 70%, 62%, and 66% of the time for the GO-MFO, GO-CCO, and GO-BPO aspects, respectively.

**Table 3:** Recall of Protein-Vec applied to held out proteins at different sequence similarity thresholds. The recall of Protein-Vec is reported for the held out 50% and held out 20% sequence similarity datasets (the held out proteins can share at most 50% and 20% sequence similarity, respectively, to any proteins in the training dataset). For each held out protein, we use Protein-Vec search to retrieve the nearest neighbor protein, and our predicted annotations are the nearest neighbor protein’s annotations. For a given held out protein, we define recall as the percentage of the predicted annotations for the aspect that are among the ground truth annotations for that aspect.

| **Aspect** | **Held Out 50 (%)** | **Held Out 20 (%)** |
| --- | --- | --- |
| EC | 87% | 61% |
| Pfam | 93% | 80% |
| Gene3D | 95% | 86% |
| GO-MFO | 80% | 60% |
| GO-CCO | 72% | 50% |
| GO-BPO | 75% | 59% |

**Table 4:** Exact match accuracy of Protein-Vec applied to held out proteins with different sequence similarity thresholds. The exact match accuracy of Protein-Vec is reported for the held out 50% and held out 20% sequence similarity datasets (the held out proteins can share at most 50% and 20% sequence similarity, respectively, to any proteins in the training dataset). For each held out protein, we use Protein-Vec search to retrieve the nearest neighbor protein, and our predicted annotations are the nearest neighbor protein’s annotations. For a given held out protein, an exact match implies that all of the predicted annotations for the aspect are identical to the ground truth annotations for that aspect (there are often multiple labels for proteins).

| Aspect | Held Out 50 (%) | Held Out 20 (%) |
| --- | --- | --- |
| EC | 87% | 61% |
| Pfam | 88% | 73% |
| Gene3D | 91% | 79% |
| GO-MFO | 70% | 48% |
| GO-CCO | 62% | 40% |
| GO-BPO | 66% | 51% |

We next compare our multi-label, multi-classification performance to new proteins that were deposited to Swiss-Prot after May 25 2022 (Table 5). Time-based splits are common in protein function prediction [21], so it naturally suits our own evaluation of Protein-Vec. Our recall predicting EC numbers (58%), Gene3D (94%), and Pfam (87%) annotations are all high on this benchmark dataset. For context, when we compare Protein-Vec with CLEAN [34], Protein-Vec achieves precision of 60% and recall of 58%, compared to CLEAN’s of 52% and 51%, when predicting enzyme commision numbers for new proteins (Table 6). As can be seen in Table 5, Protein-Vec performs better than Aspect-Vec for every single aspect on the time-based benchmark. The difference between Protein-Vec and Aspect-Vec is greatest for protein function prediction, where Protein-Vec has much higher recall than Aspect-Vec. Protein-Vec has 28%, 29% and 27% higher recall rates for GO-BPO, GO-CCO, and GO-MFO prediction, respectively. This suggests that the infusion of sequence, structure, and functional information into Protein-Vec’s model leads to better performance than any of the Aspect-Vec models on their own, which were only trained on single aspect triplets. For Pfam prediction, we compare our performance on this held-out dataset with ProtENN [46]. As ProtENN only works for individual domains, in order to make a fair comparison, we only evaluate on single domain proteins from the time-based split. On this subset of proteins, Protein-Vec has an accuracy of 90%, compared to ProtENN’s accuracy of 20%. We attribute the low performance of ProtENN to the fact that it expects domains, and we feed it single-domain proteins (with additional amino acid residues on either side of the domain).

**Table 5:** Recall of Protein-Vec and Aspect-Vec applied to new proteins for each aspect. Here, the test dataset comes from a time-based split and includes proteins deposited to Swiss-Prot after May 25th 2022 [55]. For our predictions, we used Protein-Vec and Aspect-Vec search to retrieve the nearest neighbor proteins, and our predicted annotations are the nearest neighbor protein’s annotations. For a given test protein, we define recall as the percentage of the predicted annotations for the aspect that are among the ground truth annotations for that aspect.

| Aspect | Protein-Vec | Aspect-Vec | Sample Size |
| --- | --- | --- | --- |
| GO-BPO | 42% | 14% | $n = 1131$ |
| GO-CCO | 56% | 27% | $n = 1506$ |
| GO-MFO | 58% | 21% | $n = 1296$ |
| Gene3D | 94% | 93% | $n = 1615$ |
| Enzyme Commission Number | 58% | 58% | $n = 484$ |
| Pfam | 87% | 81% | $n = 1916$ |

**Table 6:** Comparison of different Enzyme Commission Number prediction methods on a time-based split. The test dataset consists of 438 proteins deposited to Swiss-Prot after May 25th 2022 (only includes proteins shorter than 1000 amino acids to accommodate the CLEAN methods’ protein length criteria). Protein-Vec achieved the highest precision, recall, and exact match accuracy on this benchmark.

| Model | Exact match accuracy | Precision | Recall |
| --- | --- | --- | --- |
| Protein-Vec | 55% | 60% | 58% |
| Aspect-Vec (EC) | 54% | 58% | 57% |
| CLEAN | 45% | 52% | 51% |

We next compare Protein-Vec’s performance predicting GO annotations for new proteins with a group of widely used, state-of-the-art, GO annotation prediction methods. Protein-Vec outperforms these baseline methods in terms of micro-averaged and macro-averaged AUPR (Table 7). Protein-Vec even outperforms methods with access to protein structures (Table 8). Due to Protein-Vec’s mixture-of-experts model integrating 7 different aspects of protein information, our sequence representations are quite useful, as simply transferring the annotations from the nearest training set protein to the test set protein in our method’s embedding space greatly outperforms all other methods. DeepGOPlus, when compared to our method’s pipeline, includes only information from sequence and the GO labels it is trained on. DeepFRI, similarly, includes information from sequence and GO labels, but does have structure as an additional source of information. ProtTrans has the least amount of information available to it during training, since it is a self-supervised model based only on sequence, and accordingly performs less well on this task when compared to the other methods. Compared to these methods, Protein-Vec produces more informative features for protein function prediction, combining information about proteins spanning different aspects of function, enzymatic activity, protein family relationships, domain annotations, and structure similarity.

**Table 7:** Area under the Precision-Recall Curve Results for Swiss-Prot Proteins deposited after May 25th 2022. Results shown for all three branches of GO.

| Results for Test Set Swiss-Prot Proteins |  |  |  |  |  |
| --- | --- | --- | --- | --- | --- |
| Branch | Metric | Protein-Vec | deepGOPlus | prott5 | Naive |
| Molecular Function | Macro AUPR | **0.7218** | 0.5130 | 0.1402 | 0.5014 |
|  | Micro AUPR | **0.6814** | 0.1211 | 0.0174 | 0.0287 |
| Cellular Component | Macro AUPR | **0.5958** | 0.4540 | 0.1154 | 0.5025 |
|  | Micro AUPR | **0.5305** | 0.0911 | 0.0488 | 0.0113 |
| Biological Process | Macro AUPR | **0.6444** | 0.4790 | 0.1989 | 0.5008 |
|  | Micro AUPR | **0.5368** | 0.0510 | 0.0039 | 0.0036 |

**Table 8:** Area under the Precision-Recall Curve Results for those Swiss-Prot Proteins deposited after May 25th, 2022, that additionally have their structures included in the Swiss-Model database. Results shown for all three branches of GO.

| Results for Test Set Swiss-Prot Proteins with Swiss-Model Structures |  |  |  |  |  |  |
| --- | --- | --- | --- | --- | --- | --- |
| Label | Metric | Protein-Vec | deepGOPlus | DeepFRI | prott5 | Naive |
| Molecular Function | Macro AUPR | **0.7766** | 0.5252 | 0.4262 | 0.3621 | 0.3110 |
|  | Micro AUPR | **0.6610** | 0.2080 | 0.1268 | 0.0346 | 0.0037 |
| Cellular Component | Macro AUPR | **0.6327** | 0.5671 | 0.3237 | 0.3515 | 0.3991 |
|  | Micro AUPR | **0.4655** | 0.2503 | 0.1469 | 0.1034 | 0.0039 |
| Biological Process | Macro AUPR | **0.6282** | 0.5410 | 0.4339 | 0.4564 | 0.4462 |
|  | Micro AUPR | **0.5341** | 0.3901 | 0.0852 | 0.0274 | 0.0015 |

## Discussion

In this work, we have shown that Protein-Vec achieved state-of-the-art results predicting the enzyme commission numbers, Pfam annotations, and GO annotations of proteins. Through benchmarking on held-out proteins at different sequence similarity thresholds, we have demonstrated that Protein-Vec can be used to annotate remote homologs. Additionally, by showing that Protein-Vec can accurately predict enzyme commission numbers, Pfam annotations, and GO annotations of proteins in a time-based split, we have demonstrated that Protein-Vec can be deployed to annotate newly discovered proteins as well.

For each of the aspects that Protein-Vec can predict, there are different downstream applications that are enabled. By predicting enzyme commission numbers, Protein-Vec enables applications in enzyme engineering, synthetic biology and functional genomics (among many other applications). By predicting Pfam annotations, Protein-Vec enables the annotation of remote homologs and potentially aids in protein function prediction. By predicting GO annotations, Protein-Vec enables genome annotation, the functional annotation of genes, the prioritization of candidate genes (i.e. for drug discovery) among other potential applications.

In light of the impressive results of Protein-Vec, there are also several limitations to discuss. For some of the aspects that we make predictions for, like GO annotations, there are potentially many labels (10+) for proteins. The annotation scheme that we presented in this work simply involved transferring the annotations of nearest neighbor to query proteins. This annotation scheme leads to poor performance on the Aspect-Vec (GO) benchmark, and leaves room for improvement. In future work, we would like to build more sophisticated annotation transfer schemes, either employing message passing from the nearest neighbors, Gaussian processes as the CLEAN authors do [34], or another pipeline. While we demonstrated that Protein-Vec could accurately annotate remote homologs, it does not perform well on the DIAMOND benchmark (compared to DIAMOND), suggesting that Protein-Vec is not well suited to the task of local similarity, as was also the case with TM-Vec [41].

There are currently tens of billions of unannotated proteins, and the number of new sequences keeps growing at an astounding rate. As we show, Protein-Vec significantly outperforms state of the art function prediction methods, it can assist in the effort of assigning accurate functions to newly discovered proteins. This is a deeply important task, as our understanding of biology, including the mechanisms of disease, relies heavily on our knowledge of protein function.

## Online Methods

We first describe our Aspect-Vec models that can produce specific aspect-aware embeddings of protein sequences. We then use these Aspect-Vec models as experts, to build a multi-aspect-aware model (mixture of experts), which we call Protein-Vec. Using Protein-Vec, we built large searchable databases of protein representations that can be queried for finding proteins with similar sequences, structures, and functions, using only their sequence information.

### Aspect-Vec embedding model

Each Aspect-Vec model is trained on protein sequence triplets (anchor, positive, and negative examples) for the specific aspect. For every protein in a training sample (anchor protein, positive sample, negative sample), the protein sequence gets fed into the Aspect-Vec pipeline, where a pre-trained deep protein language model, ProtTrans (‘ProtT5-XL-UniRef50’)[54], first produces embeddings for every residue in each protein. Then, these residue embeddings get fed into a neural network that we train, called Aspect-Vec (Figure 1). Figure 3A shows the function which takes residue embeddings and produces a flattened vector representation for each protein for each aspect. This function, *ϕ*, is composed of several transformer encoder layers followed by average pooling and fully connected layers (see Figure 3A for details). Finally, each Aspect-Vec model produces a 512-dimensional flattened vector representation of proteins.

We used a triplet margin loss function for the Aspect-Vec models, which gets applied to the vector representations of each protein in the triplet. Figure 3A shows the triplet margin loss function that we used, which incorporates the distance between the positive and negative example labels as the margin in the triplet loss.

### Protein-Vec embedding model

The Protein-Vec model is trained on protein sequence triplets (anchor, positive, and negative examples) and the protein aspects for which they share or don’t share labels along (any combination of protein aspects we train on) (Figure 1). At every training step, we probabilistically sampled the aspects that we wanted to encode (i.e. some combination of sequence, structure, and function features). Then, we applied the corresponding Aspect-Vec model to every aspect included in the training step, producing a vector representation for each protein for each relevant aspect. Next, we combined these vectors into an Aspect-Vec matrix, and included a mask for the aspects that are not included in the training step (our model will ignore those aspects). After producing an Aspect-Vec matrix for each protein in the triplet, we applied a triplet neural network *θ* to each protein (what we call a mixture of experts model) (Figures 1, 3B). Figure 3B shows that the architecture of *θ*, which consists of several encoder layers followed by average pooling and fully connected layers. Just like the Aspect-Vec model, a Protein-Vec model produces a 512-dimensional flattened vector representation for each protein.

We used a multi-objective loss function to train Protein-Vec. The first component of the loss function is a triplet margin loss, applied to the vector representations of each protein in the triplet. Figure 3B shows the triplet margin loss function that we used, which again incorporates the distance between the positive and negative example labels as the margin in the triplet loss. When a given training sample has TM-Scores as one of its aspects, our second objective is to predict the TM-Scores between the anchor and positive example proteins, and between the anchor and negative example proteins. For the second loss component, we calculated the L1 distance between the ground truth TM-Scores and the cosine distances of the protein pairs, and add up the L1 distances for the positive and negative pairs for this component of the loss function. Figure 3B outlines this TM-Score loss component. During training, we add up the triplet margin loss and the TM-Score loss, and if a given training example does not have TM-Score as an aspect, the TM-Score loss component is set to zero.

### Protein-Vec search

We built large databases of searchable protein embeddings by starting with large databases of protein sequences. Among the different databases of protein sequences that we encoded were Swiss-Prot [55], and UniRef50 [52]. For our lookup databases, we encoded protein sequences using the relevant aspect information, and we built indexed vector-searchable database embeddings using the FAISS package [56]. When a database is queried with a new sequence, we first embed the protein using a forward pass of a Protein-Vec embedding model, and then we return the nearest neighbors of the query according to cosine similarity (the proteins in our lookup database with the highest predicted similarity to the query) (Figure 2).

Once the nearest neighbors for a query protein are returned, the outputs can easily be connected to DeepBLAST to additionally return the structural alignments for the nearest neighbors [41]. Alternatively, the outputs can be re-ranked according to another aspect that the user cares about (i.e. a user can first query along one aspect, and then re-rank the results using another aspect model).

### Protein-Vec dataset generation

We generated a dataset of protein triplets along different aspects for our training dataset, covering sequence, structure, and function aspects. All of the proteins that we trained on come from the Swiss-Prot database [55]. For every aspect, we generated a dataset of triplets where every data sample consisted of an anchor protein, a positive example and a negative example protein. For our sequence aspects, we used family and domain annotations, coming from Pfam and Gene3D. For Pfam, a positive example is defined as a protein sharing at least one Pfam annotation with the anchor protein. For Gene3D, a positive example is defined as a protein sharing at least one Gene3D annotation with the anchor protein. For negative examples, we performed hard negative mining to ensure that the classification between positive and negative examples was a hard task. For the Pfam aspect, we created hard negative samples by using Pfam clan information; as 45% of Pfam sequences belong to evolutionarily related Pfam clans, we sampled proteins with different Pfam annotations that shared at least one Pfam clan. For the Gene3D aspect hard negative mining samples, we used the Gene3D ontology to sample negative proteins that shared either a domain with the same architecture or the same topology as the anchor protein. For our structural aspect, we used TM-Score as our measure of structural similarity by running TM-align for pairs of proteins (applied to their experimental structures in the Protein Data Bank (PDB)) [33, 57] to identify structurally similar and structurally different proteins. For our functional features, we used EC number and GO information. While it is straightforward to build and sample triplets for sequence and structure features, it is less straightforward for function, as there are existing ontologies to learn across, and these ontologies have well cataloged issues including misannotation, sparsity, and over-enrichment for certain terms [39, 40]. For EC, we could leverage its compact 4 level hierarchy when we constructed negative examples. For positive examples, it was straightforward to use proteins that shared at least one EC annotation (across all 4 levels). For negative examples, we sampled proteins that did not share any EC annotations with the anchor protein but had an EC annotation that was consistent with the anchor protein’s annotation across the first 2 or the first 3 levels of the EC hierarchy (i.e. for an anchor protein with an EC annotation of 2.3.1.199, a positive sample would also have the annotation of 2.3.1.199, and the negative example could have an annotation of 2.3.X.X or 2.3.1.X). For GO, it was slightly more complicated to generate triplet, as many proteins have tens of GO annotations. For GO, we leveraged the tool GOGO [58] in order to heuristically generate triplets. We generated triplets for each GO component: the biological process, cellular component, and molecular function components. For each GO component, a positive label was considered if a pair of proteins had a high GO semantic similarity [58] and a negative label if the pair had a low GO semantic similarity. We set a GOGO threshold of .65 as being ‘similar’ for triplet generation purposes. We did not explicitly hard negative mine for GO, although we did set a minimum similarity threshold of .25 for negative examples.

As we trained on individual aspects of proteins, as well as combinations of aspects, we also sampled for protein triplets with consistent labels across multiple aspects (i.e. for a combination of aspects and a given anchor protein, the positive sample shares labels with the anchor protein across the combination of aspects, and the negative sample has a different label across the combination of aspects). As an example, for a training sample for the aspects Gene3D, Pfam, and TM-Score, the positive sample will share the same Gene3D and Pfam annotations as the anchor protein and will be structurally similar to the anchor protein (TM-Score above the similarity threshold), and the negative sample will have different Gene3D and Pfam annotations as the anchor protein and will be structurally dissimilar from the anchor (TM-Score below the similarity threshold).

The final dataset that we trained on consisted of approximately 450 million protein triplets, as well as the combination of aspects used to make each specific triplet. We further split this dataset into training, validation, and test datasets.

Our test dataset consisted of two different groups of proteins. The first group of test proteins comes from a time-based split, and consisted of proteins deposited to Swiss-Prot after May 25th 2022. The second group of test proteins are filtered for proteins that have different levels of sequence similarity to proteins that were trained on (i.e. to any anchor, positive, or negative sample proteins). We ran MMseqs2 [59] on the Swiss-Prot proteins that we studied [55], setting sequence similarity thresholds of 20%, and 50%, and sampled from the resulting clusters in order to build our 20% and 50% held-out dataset of proteins.

### Benchmark datasets

For the different benchmark datasets described below for individual aspects, we followed the same protein train/test splits of the benchmarks, and built triplet datasets for the individual aspects of each benchmark.

### Enzyme commission number benchmark

We benchmarked Protein-Vec and Aspect-Vec (EC) against CLEAN [34]. The authors of CLEAN similarly used UniProt for their training/validation/testing, and they performed a time-based evaluation where they predicted annotations for held-out enzymes from UniProt that were introduced to the database after their training cutoff date (April, 2022). In this benchmark, we evaluate the exact match accuracy, precision, and recall of CLEA, Aspect-Vec (EC) and Protein-Vec. All of these evaluations are done on the entire EC number (i.e. including the 4th and final digit in the EC hiearchy).

CLEAN, contrastive learning–enabled enzyme annotation, is another deep learning approach that learns an embedding space of enzymes by using a contrastive loss function that minimizes the distance between anchor and positive enzymes, while maximizing the distance between anchor and negative enzymes. The authors similarly employ hard negative mining to make the contrastive learning task more difficult. Once embeddings have been made using CLEAN, the method fits a two-component Gaussian mixture model on the Euclidean distances between enzyme sequence embeddings and different EC number embeddings, and the method uses this Gaussian mixture model to assign annotations and confidence intervals to predictions. Unlike Protein-Vec, Aspect-Vec (EC) and TM-Vec, they applied average pooling on the LLM they used, ESM-1b [60] right away, thereby losing any residue level information during the embedding learning process.

### Pfam benchmark

In order to evaluate our ability to perform Pfam remote homology detection, we used a benchmark dataset that was constructed by clustering Pfam-seed v32 based on sequence identity, where there was single-linkage clustering at 25% sequence identity within each Pfam family [38]. This test benchmark dataset consisted of 21,293 distant held-out Pfam sequences. This dataset was split by Bileschi et al. [46], such that every protein family was split into train, dev and test sequence sets, with a sufficient sequence distance between each set. The details of how these splits were made are detailed in the paper [46]. The metric that is reported in this benchmark is the error rate %, which refers to the % of the Pfam predictions for the held out test dataset that are incorrect (compared to the full ground truth Pfam annotations).

The different methods included in this benchmark include: BLASTp [48], phmmer [61], Top Pick HMM [61], ProtCNN [46], and ProtENN [46]. BLASTp is an alignment approach that ranks sequences by their target sequence similarity. With phmmer, test sequences are queried against a lookup database of training sequences, and the closest match is found using phmmer. The top Pick HMM is outlined in [46], and revolves around using HMMER with a top pick strategy. ProtCNN, uses a CNN architecture which first maps protein sequences to a feature vector and then applies fully connected layers to these embeddings to predict Pfam classes. ProtENN predictions are based on a majority vote from an ensemble of 19 ProtCNN models [46].

We also compared ProtENN’s performance with Protein-vec’s on single domain proteins deposited to Swiss-Prot after May 25th 2022. Perhaps because ProtENN was trained/tested on individual domains, it was not able to accurately predict Pfam annotations for single domain proteins.

### GO benchmark

To evaluate Protein-Vec on protein function prediction, we turned to a benchmark used by DeepGOPlus [16], along with different methods benchmarked in the paper. The benchmark is a time-based dataset that comes from Swiss-Prot. The training dataset consists of Swiss-Prot reviewed proteins with experimental annotations that were published before January 2016. The test dataset consists of Swiss-Prot reviewed proteins with experimental annotations that were published between January 2016 and October 2016.

The different methods that were benchmarked included: DeepGOPlus [16], which leverages sequence similarity and sequence motifs in a deep learning model that uses CNNs; DeepGO [15], a predecessor of DeepGOPlus which uses protein interaction networks as features; DiamondBLAST [49], which uses sequence similarity scores obtained by BLAST to transfer annotations; and Naive classification, which assigns GO classes to proteins based on GO annotation frequencies [21].

We also compare Protein-Vec’s performance with other function prediction methods on proteins deposited to Swiss-Prot after May 25th 2022 to ensure no metadata about test set proteins was available to our model at any point. On this benchmark, we use the latest version of the DeepGOPlus webserver (version 1.0.13), and their recommended probability threshold of 0.3. In this evaluation we also included the sequence embedding method ProtTrans [54], a transformer encoder-decoder model trained with a masked language model objective, and the Naive frequency-based method mentioned previously as baseline methods. For ProtTrans, we use the default setting in the bio-embeddings Python package that giving similarity scores of the top 3 nearest neighbors in the training set to each query protein in the test set.

In an additional evaluation that included the state-of-the-art structure-based function prediction method DeepFRI[17], we considered only those proteins that had SwissModel structures within our test set.

In these evaluations, we used the Area Under the Precision Recall (AUPR) curve to measure performance of these models. We show both macro- and micro-averaged AUPR scores for all models. In the macro-averaged score, we took each GO term separately to calculate AUPR for each term individually, and take the mean of the resulting scores. In micro-averaging, we treated the entire set of predictions and annotations as a single vector, and then computed the AUPR over the entire set of predictions at once.

### DIAMOND benchmark

The DIAMOND benchmark was introduced in the DIAMOND research paper [51]. It consists of a vast lookup dataset and a large query dataset, both comprising single and multi-domain proteins. The lookup dataset, sourced from UniRef50 (released on September 14, 2019) [52], contains 37.5 million sequences. To create a representative dataset, the authors reduced this dataset to 7.74 million representative protein sequences along with their SCOP family annotations [53]. The query dataset, obtained from the NCBI nr database (released on October 25, 2019), also incorporates SCOP family annotations for proteins. The authors confined this dataset to a maximum of 1,000 protein sequences for each SCOP superfamily, resulting in a dataset of 1.71 million queries.

## Code availability

Protein-Vec can be found on GitHub at: https://github.com/tymor22/protein-vec.

## Acknowledgements

This research was supported by NIH R01DK103358, the Simons Foundation, NSF IOS-1546218, R35GM122515, NSF CBET-1728858 and NIH R01AI130945 to T.H.; the Flatiron Institute as part of the Simons Foundation to R.B.; and the Samsung Advanced Institute of Technology (Next Generation Deep Learning: from pattern recognition to AI), Samsung Research (Improving Deep Learning using Latent Structure) and NSF Award 1922658 to K.C.

We would like to thank the Flatiron Institute, including N. Carriero and D. Simon, for providing the computer support required to train these models and for all of their help.

